# Deep learning enables structured illumination microscopy with low light levels and enhanced speed

**DOI:** 10.1101/866822

**Authors:** Luhong Jin, Bei Liu, Fenqiang Zhao, Stephen Hahn, Bowei Dong, Ruiyan Song, Tim Elston, Yingke Xu, Klaus M. Hahn

## Abstract

Using deep learning to augment structured illumination microscopy (SIM), we obtained a fivefold reduction in the number of raw images required for super-resolution SIM, and generated images under extreme low light conditions (100X fewer photons). We validated the performance of deep neural networks on different cellular structures and achieved multi-color, live-cell super-resolution imaging with greatly reduced photobleaching.

## Main

Structured illumination microscopy (SIM) is a widely used super-resolution technique that has had substantial impact because of its ability to double resolution beyond the light diffraction limit while using wide field illumination and maintaining compatibility with a wide range of fluorophores. SIM applies varying, nonuniform illumination on samples and then uses dedicated computational algorithms to derive super-resolution information from nine or fifteen sequentially acquired images, for 2D or 3D imaging respectively. Since it was first introduced by Mats Gustafsson in 2000 ^1^, SIM has been evolving constantly to improve speed, resolution and to decrease the required light dosages. Reconstruction algorithms have been developed to estimate microscope parameters robustly^2^, minimize reconstruction artifacts^3,4^, reduce the required number of raw images^5^, and check the quality of the raw data and reconstruction^6^. The primary limitation of SIM is the need to obtain a series of high-quality images for each reconstructed high-resolution SIM image; this decreases temporal resolution and increases photobleaching.

Recently, there has been an explosion of deep learning applications in many aspects of biological research. For microscopy, deep learning has demonstrated impressive capabilities in cell segmentation/tracking, morphology analysis, denoising, single molecule detection/tracking, and super-resolution imaging. ^7–9^ Use of deep learning for content-aware image restoration (CARE) has shown great promise in de-noising, enhancing signal to noise ratio, and isotropic imaging^10^. However, the potential of deep learning in SIM has not been explored. We sought to apply deep learning to increase the speed of SIM by reducing the number of raw images, and to retrieve super-resolution information from low light samples. We accomplished this by reconstructing images using deep neural networks that had been trained on real images, enabling us to visualize specific complex cellular structures (mitochondria, actin networks etc.) and address complicated cellular or instrument-dependent backgrounds (e.g. out of focus light). The approach was named DL-SIM (deep learning assisted SIM). U-Net is one of the most popular convolutional neural network architectures^11,12^. We show that U-Net can be trained to directly reconstruct super-resolution images from SIM raw data using fewer raw images.. Where conventional SIM typically required nine or fifteen images for reconstruction, U-Net achieved comparable resolution with only three images. Also, using stacked U-Nets, we could restore high quality, high resolution images from image sequences acquired using greatly reduced light dosages. We demonstrated the capabilities of DL-SIM in multicolor super-resolution imaging of living cells.

We first tested the ability of U-Net to reconstruct super-resolution images from SIM raw sequences. Conventional SIM excites specimens with sinusoidal waves at different angles and phases. Typically, it requires 9-images (3 angles, 3 phases) for two-beam SIM and 15-images (3 angles, 5 phases) for three-beam SIM^13^. We trained a single U-Net (U-Net-SIM15) by taking 15 SIM raw images as the input and the corresponding conventional SIM reconstruction results as the ground truth (Fig. 1a, Supplementary Fig. 1, Methods). U-Net-SIM15 was trained on four different subcellular structures: microtubules, adhesions, mitochondria and F-actin. Each dataset was randomly separated into subsets for training, validation and testing respectively (Supplementary Table 1). We tested performance on cells that had not been seen by the networks during the training step. U-Net-SIM15 consistently produced images with fidelity comparable to that of the conventional SIM algorithm (Fig. 1b, column U-Net-SIM15). We next tested whether we could accelerate SIM imaging by reducing the number of raw images for SIM reconstruction. We trained another U-Net (U-Net-SIM3) using only three SIM raw images (the first phase at each angle) as input, and again used the SIM reconstruction results from fifteen images as the ground truth. Surprisingly, U-Net-SIM3 could produce restored images (Fig. 1b, column U-Net-SIM3) with the quality of those produced using U-Net-SIM15 and the ground truth. We estimated the restoration error maps using SQUIRREL^14^ for both conventional SIM reconstruction and U-Net output images against the average projection of the SIM raw data, and used both resolution-scaled error (RSE) and resolution-scaled Pearson coefficient (RSP) to quantify the quality of the restoration (Supplementary Fig. 2, Methods). Line profiles along microtubules, adhesions, mitochondria and F-actin showed that U-Net reached a lateral resolution comparable to conventional SIM reconstruction (Fig. 1b). We also quantified the resolution of each approach using a recently reported approach based on decorrelation analysis ^15^(Fig. 1c), peak signal-to-noise ratio (PSNR), the normalized root-mean-square error (NRMSE) and the structural similarity index (SSIM) (Supplementary Table 2, Methods). Furthermore, we applied the pre-trained model to visualize the dynamics of microtubules in living cells with high resolution (Supplementary Video 1).

**Figure 1:**
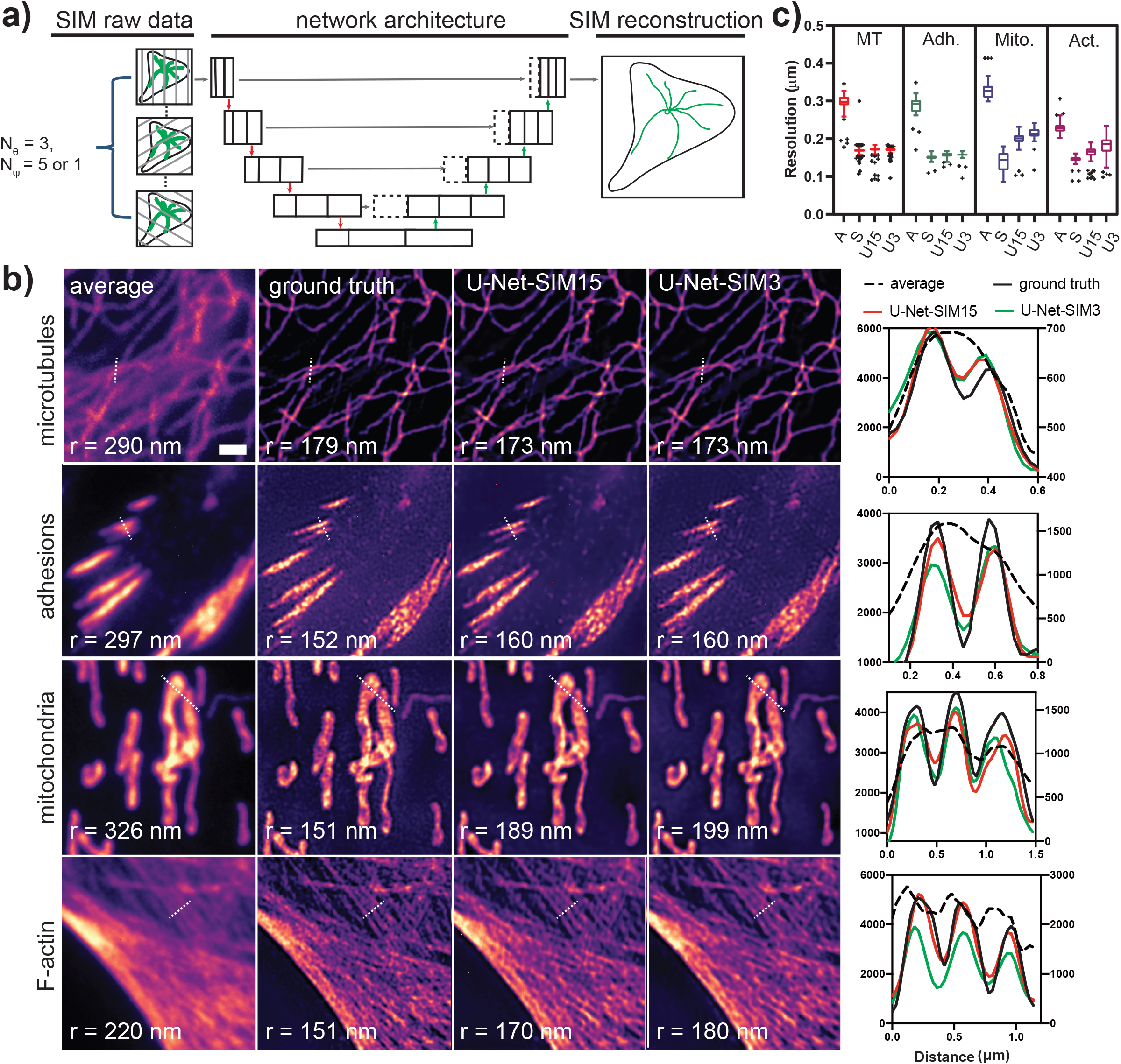
Super-resolution imaging with U-Net. **a)** Fifteen or three SIM raw data images were used as input and the corresponding SIM reconstructions from 15 images were used as the ground truth to train the U-Net. Θ: the angle of the sinusoidal patterned illumination; ψ: the phase of the patterned illumination. **b**) Reconstruction results for different subcellular structures. Shown are average projections of 15 SIM raw data images (first column), the reconstruction results from a conventional SIM reconstruction algorithm (second column), U-Net-SIM15 output (third column), U-Net-SIM3 output (fourth column) and line profiles along the dashed line in each image (fifth column). In the line profile plot, “average” is shown on the right y-axis and all others share the left y-axis. **r** indicates the resolution. **c**) The achieved resolution of different approaches was estimated. MT: microtubules; Adh.: adhesions; Mito.: mitochondria; Act.: F-actin. A: average; S: SIM reconstruction; U15: U-Net-SIM15; U3: U-Net-SIM3.

Next, we sought to increase acquisition speed and reduce photobleaching by using low laser power (10-fold reduction) and reducing the exposure time (20-fold reduction). Reducing light levels degrades conventional SIM reconstruction because there is less information in the raw data. U-Net and other machine learning methods have been successfully adopted to recover information from low light samples^10^. We therefore trained another U-Net (U-Net-SNR) to recover signals from poor quality images), and fed the output from this to the pre-trained U-Net-SIM15. U-Net-SNR alone could recover information from low light samples (e.g. periodic illumination patterns, Supplementary Fig. 3a, 3b). Combining the two networks produced good resolution throughout much of the images, but failed in some specific areas (Supplementary Fig. 3c). We hypothesized that training a deeper network by connecting two U-Nets could improve performance^10,16^, so we constructed a new architecture by chaining two U-Nets through skip-layer connections (Fig. 2a, Supplementary Fig. 4). This network (scU-Net) was trained using an input of 15 SIM raw data images obtained under low light conditions, and using SIM reconstruction from normal light dosages as the ground truth. We found that scU-Net produced better restoration quality than either SIM reconstruction or U-Net-15 (Fig. 2b) and provided the least restoration errors (Supplementary Fig. 5). We quantified network performance by calculating PSNR, NRMSE and SSIM of the different reconstruction approaches relative to the ground truth. The scU-Net achieved considerable improvements over other approaches (Supplementary Table 3). We further tested the pre-trained scU-Net to visualize the dynamics of microtubules in living cells under extreme low light conditions (Fig. 3a, Methods). The quality of the input and conventional SIM reconstruction was poor under these conditions (Fig. 3a, column 1 and 2). U-Net-SIM15 improved the reconstruction but missed some details (Fig. 3a, column 3). scU-Net produced the best results (Fig. 3a, column 4), enabling us to track the dynamics of single microtubules (Fig. 3b, Supplementary Video 2) with substantially reduced photobleaching (Supplementary Fig. 6). We used the pretrained scU-Net to examine microtubule/mitochondrial interactions with short exposure time and low laser intensity (Fig.3c, Supplementary Video 3, Methods), but with no discernable compromise to image quality.

**Figure 2:**
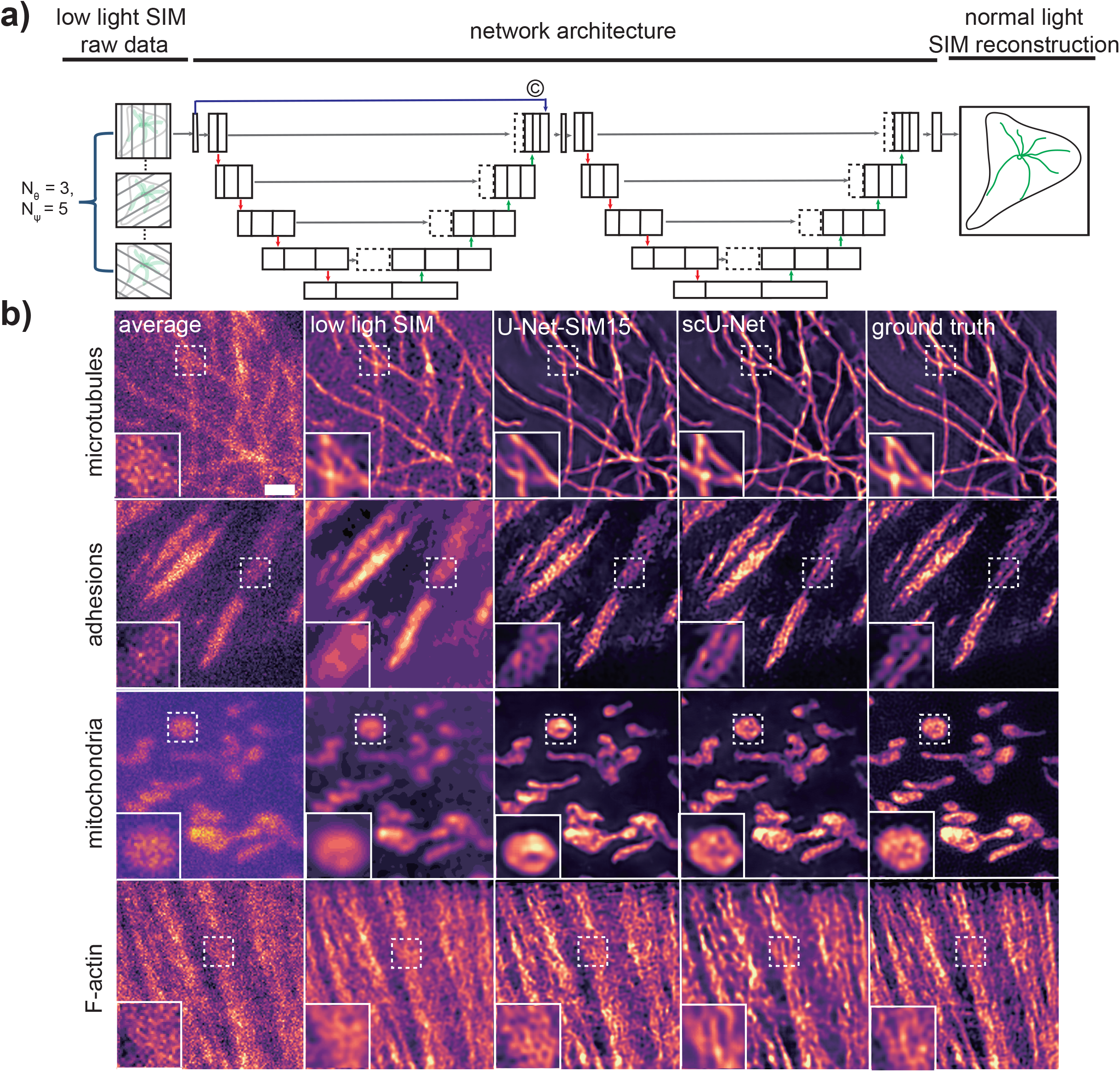
Super-resolution imaging under extreme low light conditions. **a)** Two U-Nets were stacked through skip-layer connections. Fifteen SIM raw data images taken under low light conditions were used as the input and the corresponding SIM reconstructions under normal light conditions were used as the ground truth to train the scU-Net. **b)** Reconstruction results for different subcellular structures (first row: microtubules; second row: adhesions; third row: mitochondria; fourth row: F-actin). Shown are average projections of 15 SIM raw data (first column), the reconstruction results from a conventional SIM reconstruction algorithm (second column), U-Net-SIM15 output (third column), scU-Net output (fourth column), and the ground truth from SIM reconstruction under normal light conditions (fifth column). The local enlargements show the restoration quality.

**Figure 3:**
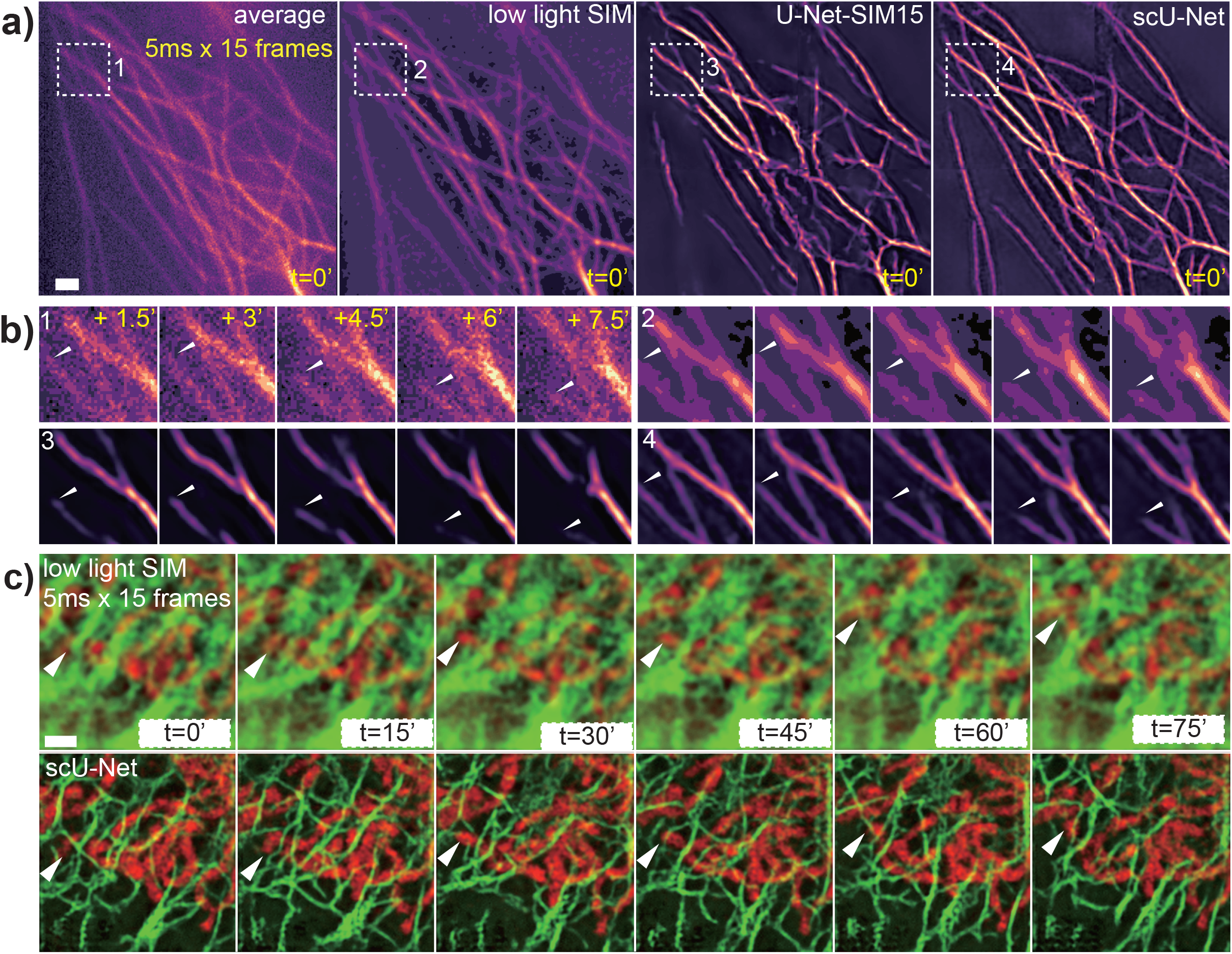
scU-Net for live cell imaging. **a**) Reconstruction results for microtubules in living cells. A representative time point is shown. First panel shows the average projections of 15 SIM raw data images; second panel shows the SIM reconstruction; third panel shows the U-Net-SIM15 output; fourth panel shows the scU-Net output. **b**) Enlarged views of areas indicated by the white-dashed box in **a** are shown. The dynamics of a single microtubule (white triangle) was well restored by scU-Net. **c**) scU-Net reveals the dynamics of microtubule-mitochondria interactions. First row: SIM reconstruction; second row: scU-Net output.

Using networks trained on synthetic tubular and point-like data, previous studies showed that U-Net could resolve sub-diffraction structures with at least 20X faster speed than super-resolution radial fluctuations (SRRF)^10,17^. We investigated the performance of U-Net trained on real biological samples. With a single U-Net (U-Net-SRRF5), we achieved comparable quality by taking as few as five total internal reflection fluorescence microscopy (TIRFM) images, 40X less images than SRRF (Supplementary Fig. 7a, 7b, 7c). In general, 50-70 single cells were needed for the training step to obtain a working model (Supplementary Table 1). A model trained from a given intracellular structure produced significant artifacts when used to examine other structures (Supplementary Fig. 8). We applied transfer learning^18^ to minimize the effort when adapting our pre-trained networks to other structures (Methods). U-Net-SIM15 pre-trained on microtubules was used to initialize a new U-Net retrained with other structures (adhesions, mitochondria and F-actin) (Methods). This achieved considerable improvement in the output quality even with only 200 training samples for each structure and 1/10 the training effort (Supplementary Fig. 9). Although we could achieve ultra-short exposure times (< 5 ms) for single frames, the minimum time interval between each processed image (1 second) was limited by our commercial SIM system, due to the time required to change the hardware between each image acquisition. Our data show that deep learning can substantially push this speed limit using home-built SIM systems^4,19^ or faster commercial systems.

## Online methods

### Microscopes

SIM imaging was performed on the Nikon N-SIM system equipped with 488 nm (70 mW), 561 nm (70 mW) and 647 nm (125 mW) laser lines, two EMCCD cameras (Andor iXon3) and a 100X, NA 1.45 objective (Nikon, CFI Apochromat TIRF 100XC Oil). For the training datasets, we used 10% intensity of 488/561/647 nm lasers and 200 ms exposure time to acquire SIM images with high quality, and 1% intensity of 488/561/647 nm lasers and 20 ms exposure time to acquire SIM images under low light conditions. For the live cell microtubule imaging experiments, we used 10% intensity of the 561 nm laser with 100 ms exposure for normal light conditions, and used 1% intensity of the 561 nm laser and 5 ms exposure for low light conditions. For the dual-color experiments, we used the 488 nm laser at 2% power and the 561 nm laser at 1% power with 50 ms exposure time to visualize microtubule-mitochondria interactions.

The SRRF imaging was performed on a home-built total internal reflection fluorescence (TIRF) microscope equipped with 488 nm, 561 nm and 647 nm laser lines, two sCMOS cameras (Photometrics, Prime 95B) and a 100 X, NA 1.49 TIRF objective (Olympus, UAPON100XOTIRF). For imaging microtubules, we used 0.6 mW from a 647 nm laser and 100 ms exposure time.

### Cell culture and preparation

For the training step, fixed cell samples were used. We prepared four training datasets from various subcellular structures, including microtubules, adhesions, mitochondria and F-actin. For imaging mitochondria and F-actin, fluorescently prelabeled commercial slides (Molecular Probes, F36924) were used. Microtubules and adhesions samples were prepared as follows:

#### Microtubules

Mouse embryonic fibroblast (MEF) cells were fixed in (−20 ℃ 100% methanol for 3min. The cells were washed for five times with 0.1% Triton-X100 in phosphate-buffered saline (PBS), and then permeabilized with 0.5% Triton-X100 in PBS for 10 min. Next, the cells were washed three times again and blocked for 15 min with blocking buffer (5% BSA in the wash buffer). The cells were incubated with beta-tubulin antibody (Developmental Studies Hybridoma Bank, E7) (1:500 dilution into blocking buffer), followed by the Alexa-647 dye (CST, 4414S).

#### Adhesions

MEF cells were fixed with 37 ℃ 4% paraformaldehyde for 10 min at room temperature. The cells were washed with PBS three times and blocked with 3% BSA, 0.2%Triton-X100 in PBS for 30min. The cells were incubated with 1:100 diluted primary antibody (Santa Cruz Biotechnology, sc-365379) for 30 min and washed (0.2% BSA, 0.1% Triton-X100) five times, followed by the staining of the Anti-rabbit IgG (H+L), F(ab')2 Fragment conjugated with Alexa-647 dye (CST, 4414S).

#### Live cells labelling

COS-7 cells were cultured in Dulbecco’s modified Eagle’s medium (Fisher Scientific, 15-013-CV) supplemented with 10% fetal bovine serum (Gemini, 100-106) and 5% GlutaMax (Thermo Fisher, 35050061). To visualize microtubules, COS-7 cells were transfected with a microtubule-associated protein (EMTB-3xmCherry or EMTB-3xEGFP), using Viromer Red transfection reagent (OriGene Technologies, TT1003102). Mitochondria were stained with MitoTracker Green FM (Thermo Fisher, M7514) according to the manufacturer’s instructions.

### Data preprocessing

For deep learning, the size of the training dataset should be as large as possible to cover the distribution of images in the task domain. However, collecting huge amount of single cell data can be time-consuming and expensive. We therefore cropped the original image stacks into smaller patches to generate more training samples for all the experiments. In SIM experiments, the size of the raw image stack was 512 × 512 × 15 (width × height × frame). To prepare the input for U-Net-SIM15, the raw stack was cropped into 128 × 128 × 15 patches. For U-Net-SIM3, only the first phase of three illumination angles were selected, producing 128 × 128 × 3 patches. After that, we manually discarded the patches, which contained only background information. We then located the same areas on the SIM reconstruction images to produce the corresponding ground-truth images. In total, we obtained 800-1500 samples for different structures, which were then randomly divided into training, validation and testing subsets. Detailed information about each dataset is in Supplementary Table 1.

Normalization is also important to the efficiency and robustness of the network. We normalized the input images to the maximum intensity of the whole input dataset and the ground truth images to the maximum intensity of the SIM reconstruction dataset.

Since U-Net requires the width and height of the input images to match the ground truth images, we resized the input dataset to 256 × 256 × C (C means the number of channels for the input data; it differs among different experiments) using bi-cubic interpolation.

### Network architectures and training details

U-Net-SIM15 and U-Net-SIM3, U-Net-SNR, and U-Net-SRRF share similar network architectures (Supplementary Fig. 1) and they only differ in the numbers of channels of either input or output (ground truth) dataset (U-Net-SIM15: C_in_ = 15, C_out_ = 1; U-Net-SIM3: C_in_ = 3, C_out_ = 1; U-Net-SNR: C_in_ = 15, C_out_ = 15; U-Net-SRRF: C_in_ = 5, C_out_ = 1; C_in_ and C_out_ are the numbers of channels of the input and output, respectively). For the U-Net-SNR, we took fifteen SIM raw data images acquired under low light conditions as the input and the same sample under normal light conditions as the ground truth. For the U-Net-SRRF, we used five TIRF frames as the input and the SRRF reconstruction from 200 frames as the ground truth. For the SIM experiment under low light conditions, the scU-Net was used (Supplementary Fig. 4). The training details for each experiment are listed in Supplementary Table 1. The loss function for all experiments is defined as:

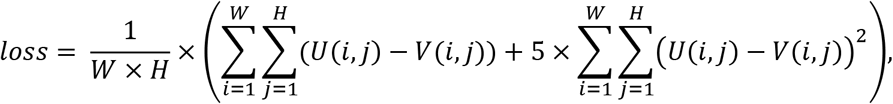

Here, ***W*** and ***H*** represent the width and height of the ground truth image in the training step (W=256, H=256 across all networks related to SIM and W=320, H=320 for the SRRF experiment). ***U*** and ***V*** represent the ground truth image and the output from the network, respectively. The codes for training and testing was written using Python with PyTorch framework. All the source codes will be available online (https://github.com/drbeiliu/DeepLearning).

### Quantification of performance for each network

For the testing part, we used four metrics to evaluate the performance of our networks, including image resolution, peak signal-to-noise ratio (PSNR), normalized root-mean-square error (NRMSE) and structural similarity index (SSIM). The resolution of each cropped image was estimated using the ImageDecorrleationAnalysis plugin in Fiji/ImageJ with the default parameter settings^15^. Note that for low light images, the image quality was so poor that the plugin failed to report a reasonable value. In that case, we used the whole-cell image, instead of the cropped patches to estimate the resolution. As for PSNR, NRMSE and SSIM, we used the SIM reconstruction results under normal light conditions as the ground truth. Each metric was calculated as below:

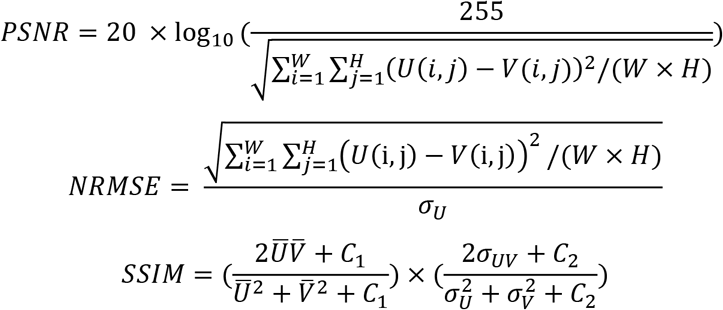

Here, ***W*** and ***H*** represent the width and height of the ground truth image in the training step (W=256, H=256 across all networks). ***U*** and ***V*** represent the ground truth image and the output of the network, respectively. 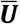 and 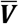 represent the averages of ***U*** and ***V***. σ_*U*_and σ_*V*_ represent the variances of ***U*** and ***V***. σ_*UV*_ is the covariance of ***U*** and ***V***. The items ***C***_**1**_ and ***C***_**2**_ are small positive constants that stabilize each term (*C*_1_ = (*k*_1_*L*)^2^, *C*_2_ = (*k*_2_*L*)^2^, ***L*** is the dynamic range of the pixel-values, ***k***_1_ = 0.01 and ***k***_2_ = 0.03 by default). The code for calculating the performance was written with Python.

We then computed the performance of each metric for each architecture based on the output of the networks and the ground truth images (Supplementary Table 2, Supplementary Table 3). RSP and RSE were introduced before to assess the quality of super-resolution data^14^ and were calculated using NanoJ-NanoJ-SQUIRREL (https://bitbucket.org/rhenriqueslab/nanoj-squirrel/wiki/Home).

### Transfer learning

Directly applying a model trained on one specific structure to other structures may produce significant artifacts (**Supplementary Fig. 8**), which means that each target needs a unique model. In theory, we need to prepare ~1000 training samples and train the network for 2-3 days (~2000 epochs) on a consumer-level graphics card (NVIDIA GTX-1080 GPU) to get a working model for each structure we tested. We adopted transfer learning^18^ to reduce the effort of imaging new structures. Briefly, we took the parameters obtained from a pre-trained network to initialize a new network and started retraining on a different structure with smaller training samples size (two hundred of cropped patches). We validated the effectiveness of transfer learning in restoring different structures. Even with reduced training efforts (200 epochs), the new model produced results comparable to the model trained with a much larger dataset and greater training effort. (**Supplementary Fig. 9**)

### SRRF experiment

In the SRRF experiment, the original input images were cropped into 64 × 64 × 5 (width × height × frame) and the original ground truth images, which were computed from 200 TIRF images, were cropped into 320 × 320 × 1. Since the size of the SRRF super-resolution image is larger than the input, we resized the cropped input image (64 × 64 ×5) into 320 × 320 × 5 using bicubic interpolation to match the size of the ground truth.

## Acknowledgments

This work was funded by the NIH (R35GM122596 to KMH), the National Key Research and Development Program of China (SQ2018YFE011598 and 2016YFF0101406 to Y.X.) and the National Natural Science Foundation of China (31571480 and 61427818 to Y.X.). We thank Tony Perdue from the Department of Biology Microscopy Core at UNC for assistance with SIM imaging.

## Author contributions

BL, LJ and KMH conceived of the project. LJ and BL performed the imaging. LJ and FZ developed the code with the help of BD, RS and SH. LJ prepared the data and figures with help of all authors. The manuscript was written with input from all authors, starting from a draft provided by BL. TE provided oversight of mathematical aspects of the project, and YKX provided insight into imaging and microscopy. BL and KMH supervised the project.

## Competing Interests statement

The authors declare no competing interests.

---

**Supplementary Figure 1.**
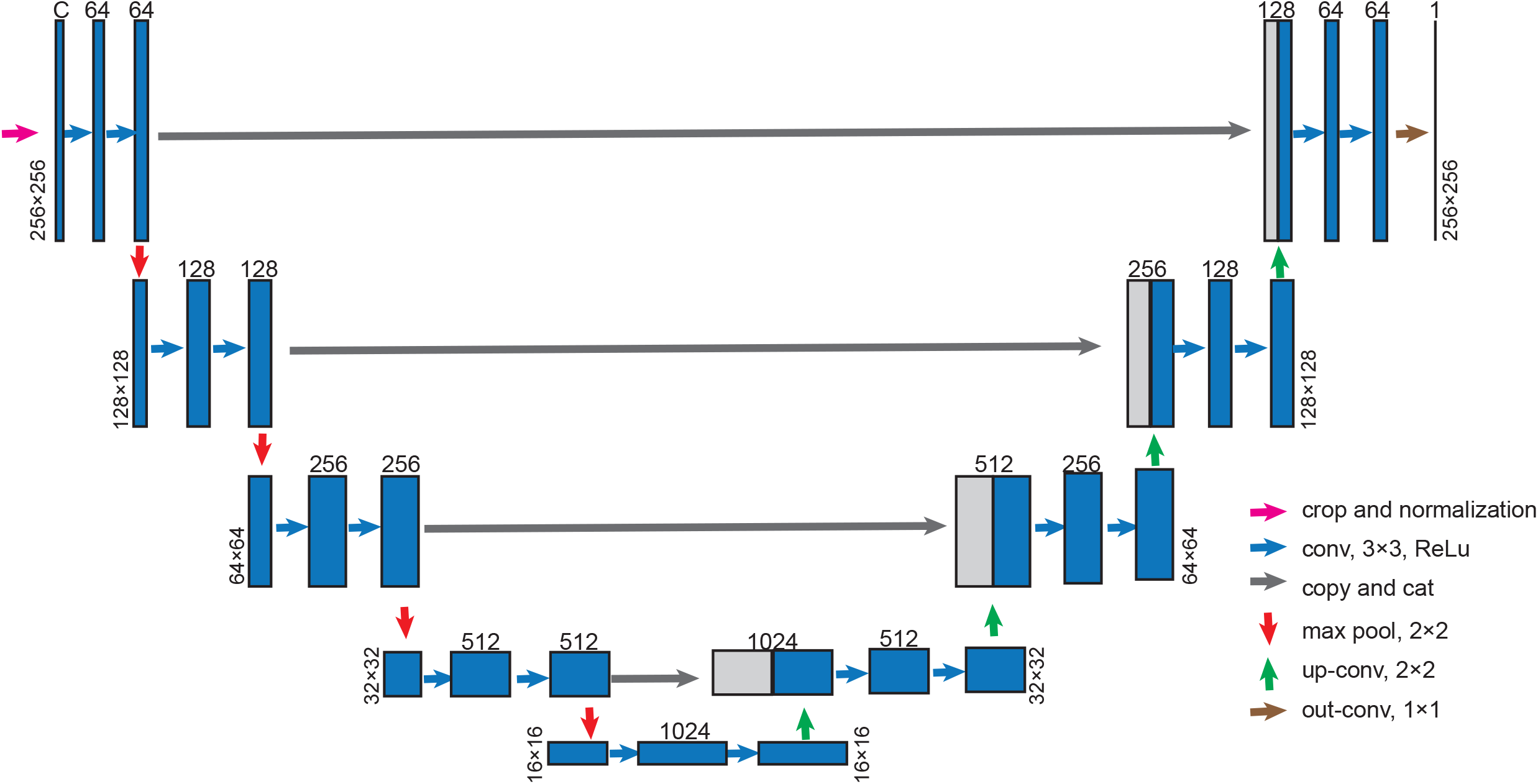
U-Net-SIM architecture. **C** indicates the channel number of the input, which differs among different networks. U-Net-SIM15: C = 15; U-Net-SIM3: C = 3.

**Supplementary Figure 2.**
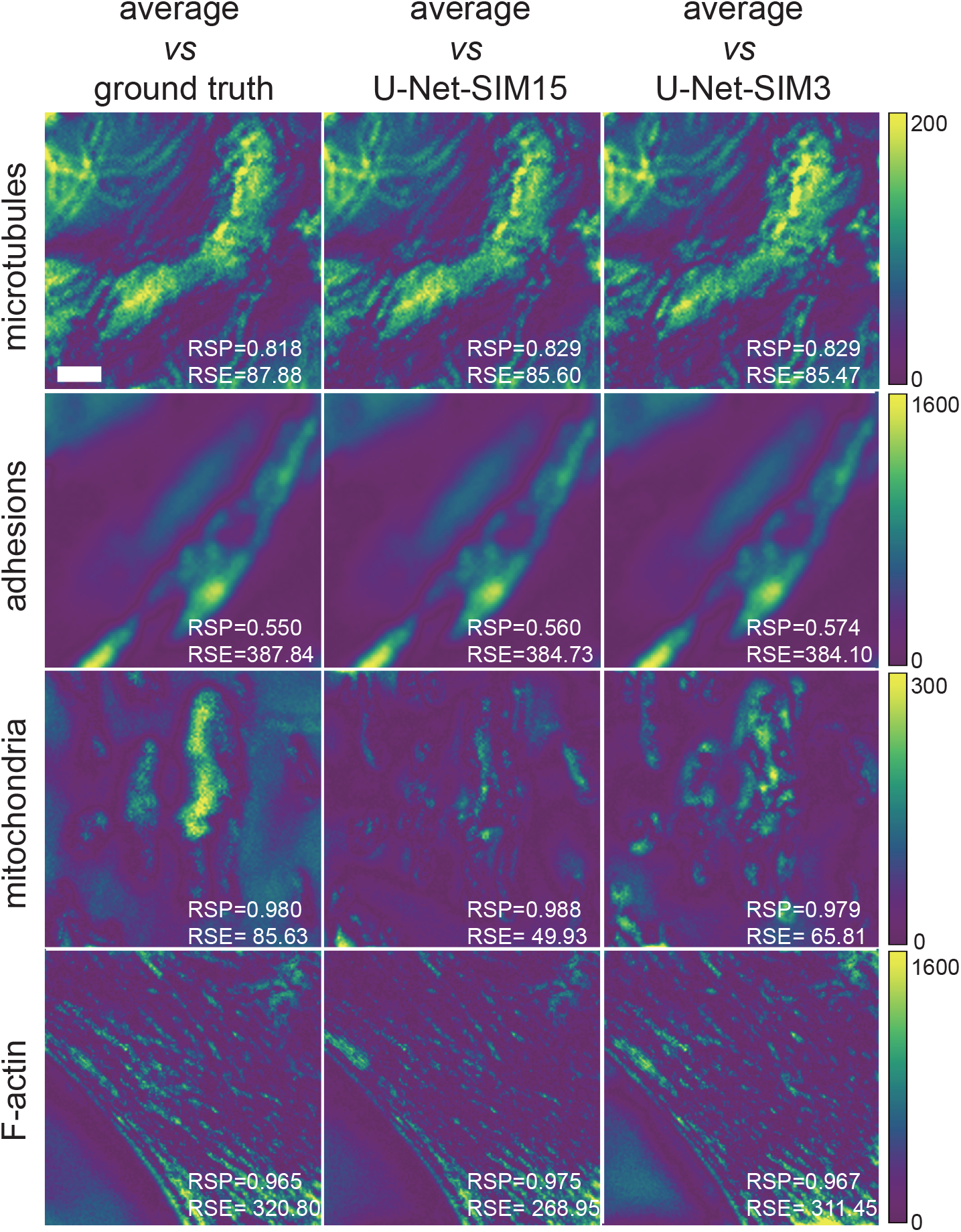
Restoration error estimation of U-Net-SIM. The error maps were estimated via SQUIRREL for SIM reconstruction, U-Net-SIM15 and U-Net-SIM3 output against the average projection of the SIM raw data. Shown are the different structures addressed in Figure 1. Two metrics were calculated to quantify the restoration: the resolution-scaled error (RSE) and the resolution-scaled Pearson coefficient (RSP). Scale bar: 1 μm.

**Supplementary Figure 3.**
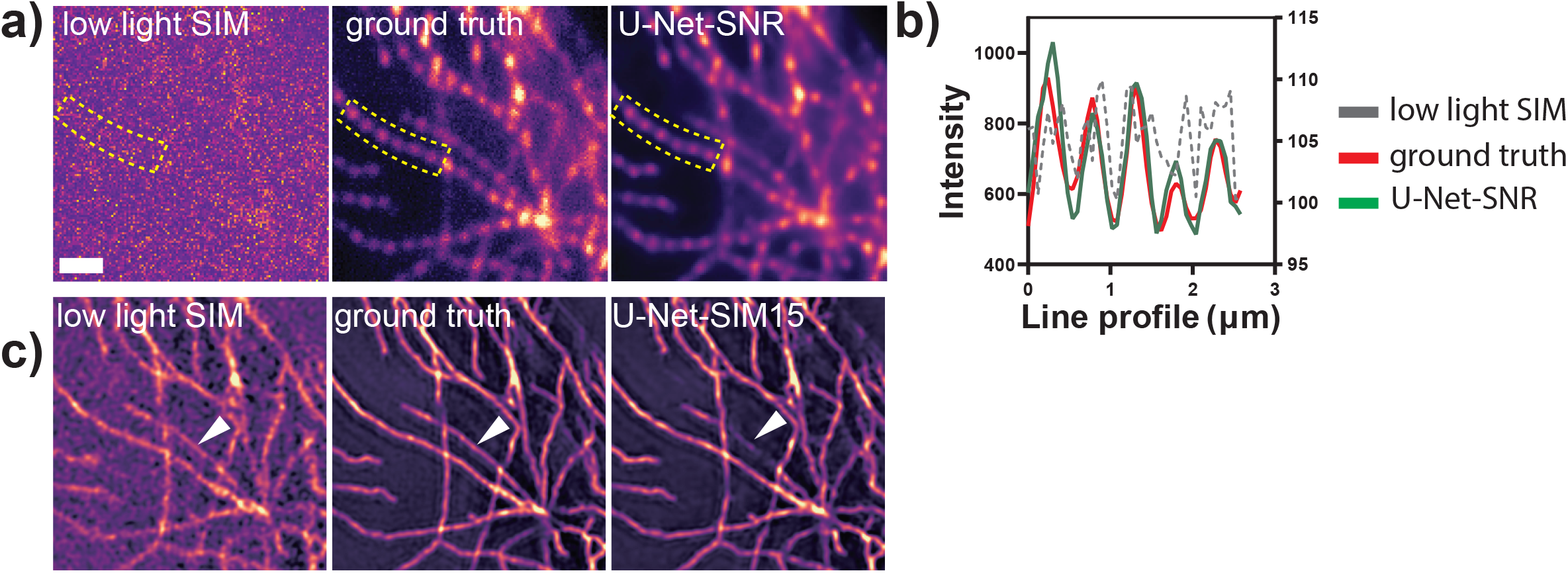
Image recovery with U-Net-SNR and the pre-trained U-Net-SIM15. **a, b)** Microtubule samples were illuminated with structured patterns under low-light (**a**, left) and normal-light conditions (**a**, middle). U-Net-SNR were able to restore the structures (**a**, right), as well as the periodic illumination patterns along single microtubule as shown in **b**. In the line profile plot, low light SIM is shown on the right y-axis and all others share the left y-axis. One out of fifteen images are shown. **c**) The output of U-Net-SNR were taken as the input of U-Net-SIM15 to retrieve SR information. Shown are SIM reconstruction results under low light conditions in the left panel of **c**, SIM reconstruction results under normal light conditions in the middle panel of c, and U-Net-SIM15 output taking images in the right panel of **a** as input. The combination of U-Net-SNR and U-Net-SIM15 achieves moderate restoration performance, but may fail at challenging areas, indicated by the white triangles.

**Supplementary Figure 4.**
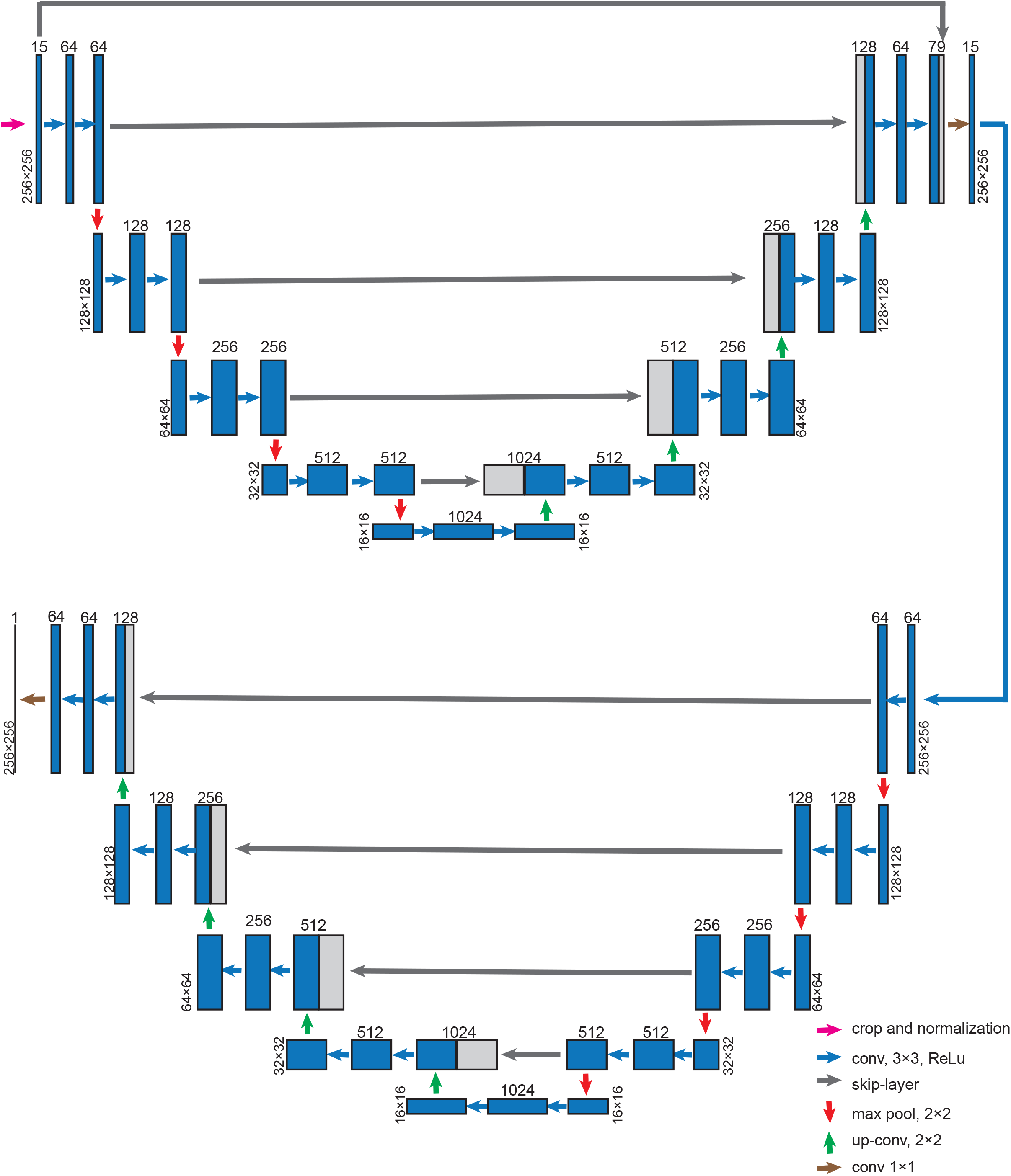
scU-Net architecture. Two U-Nets were stacked together via skip-layer connection. The network was trained by taking 15 SIM raw data images under low light conditions as the input and the SIM reconstruction under normal light conditions as the ground truth.

**Supplementary Figure 5.**
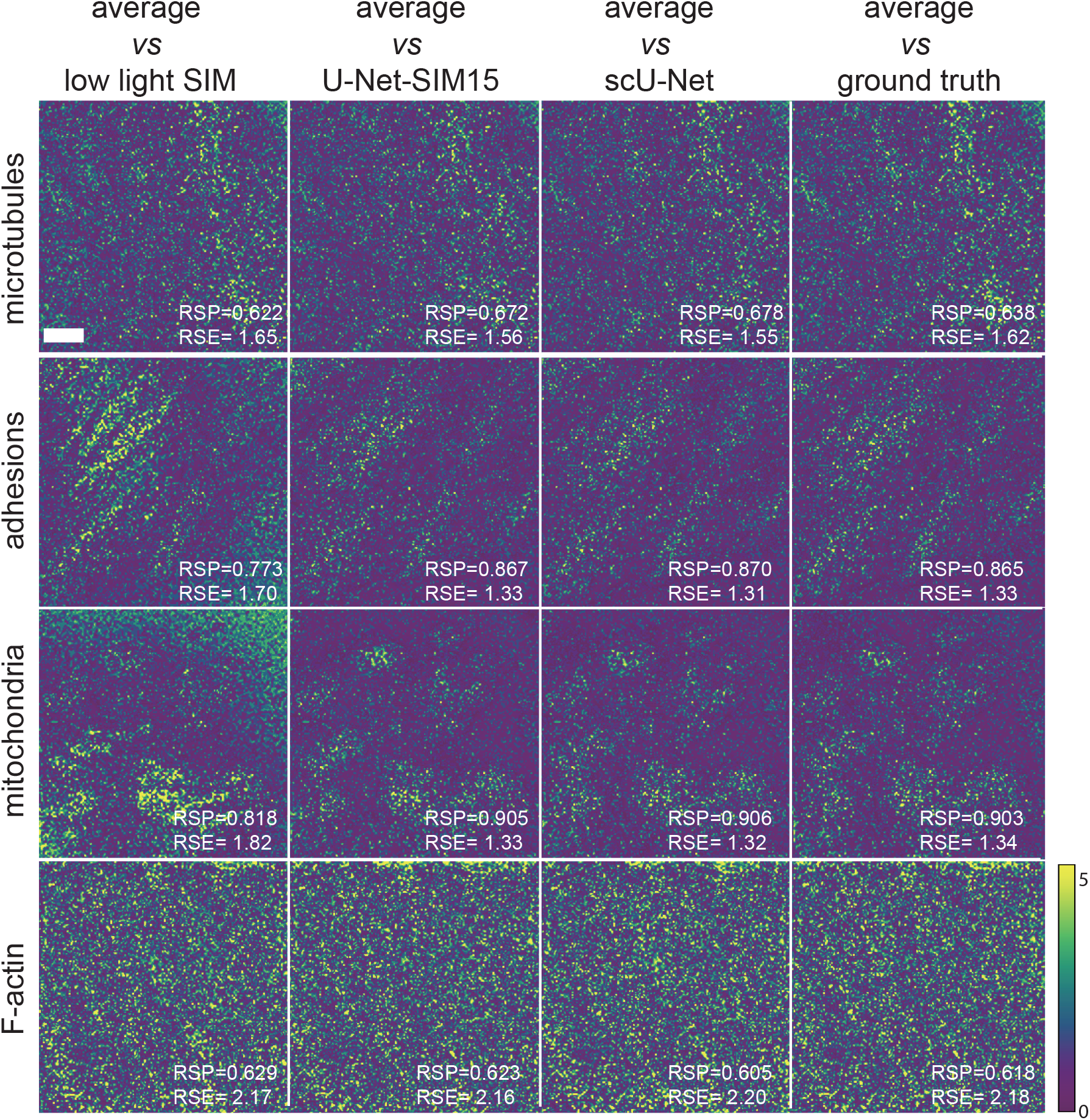
Restoration error estimation of scU-Net. The error maps were estimated via SQUIRREL for the indicated restoration approaches. Shown are the different structures addressed in Figure 2. The values of RSE and RSP were calculated to quantify the restoration. Scale bar: 1 μm.

**Supplementary Figure 6.**
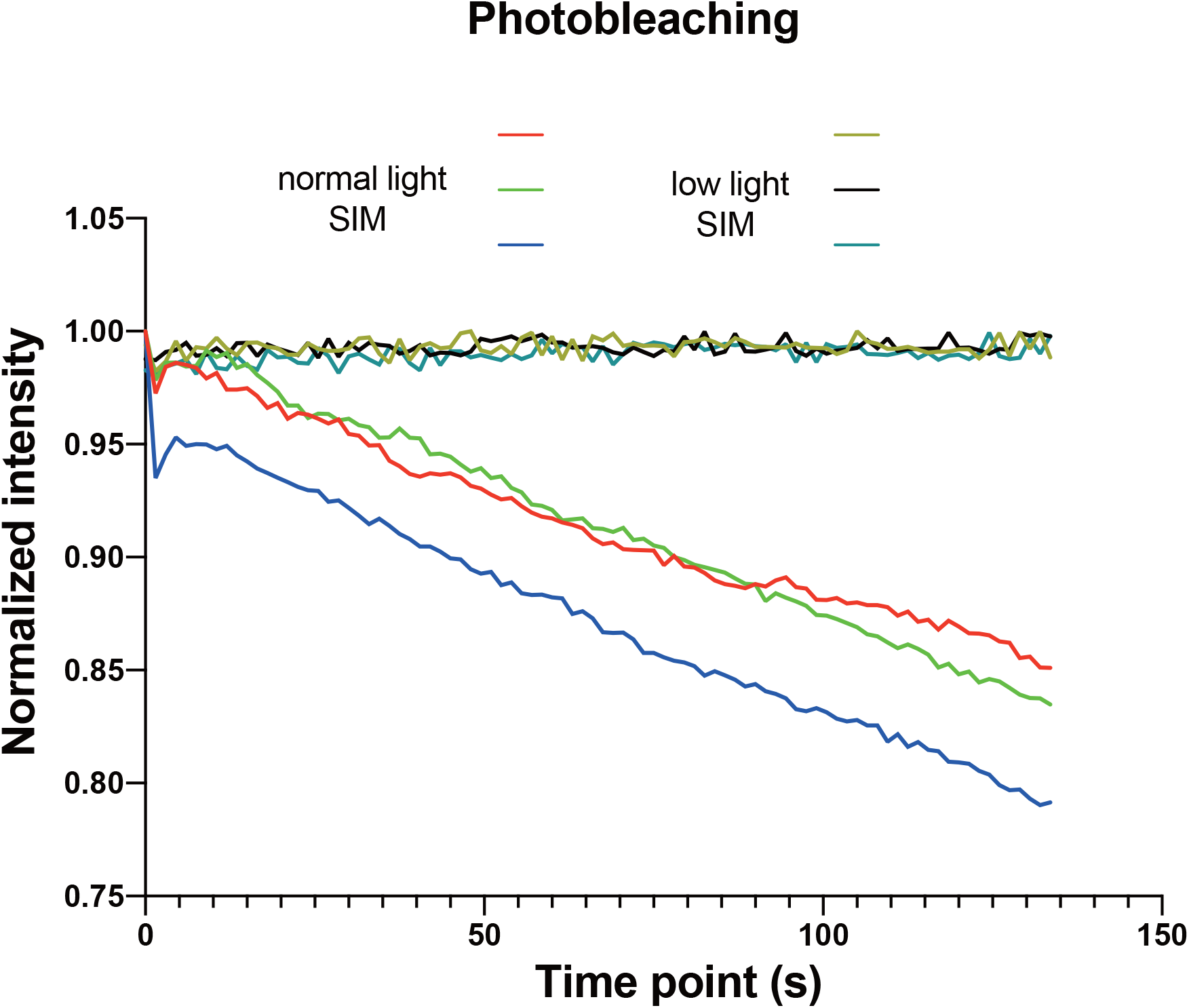
Photobleaching of SIM imaging under low light and normal light conditions. COS-7 cells were transiently transfected with EMTB-3XmCherry. For normal light SIM imaging, we used 10% 561 nm laser power and 100 ms exposure time, while for the low light SIM imaging, we used 1% 561 nm laser power and 10 ms exposure time.

**Supplementary Figure 7.**
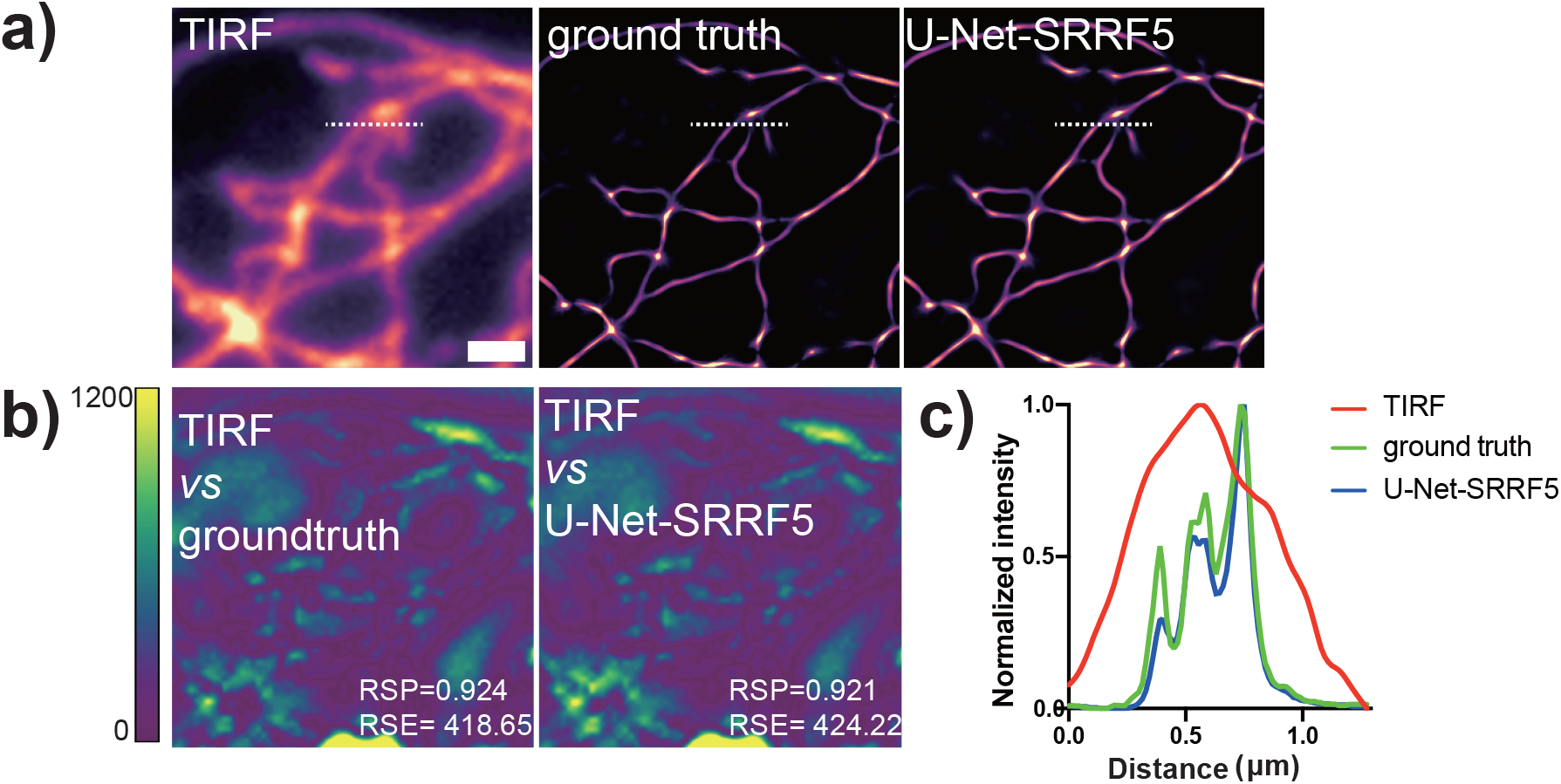
U-Net-SRRF5 restoration. MEF cells were fixed and the microtubules were stained with Alexa-647 dye. The cells were examined under a TIRF microscope. **a)** Shown are: a representative TIRF image of microtubules (**left**), SRRF reconstruction of 200 TIRF images, output of U-Net-SRRF5 by taking five TIRF images as input. **b**) Restoration error of SRRF and U-Net-SRRF. **c**) Line profiles along dotted lines in **a** show resolution improvement of U-Net-SRRF5. The value of RSE and RSP were calculated to quantify the restoration. Scale bar: 1 μm.

**Supplementary Figure 8.**
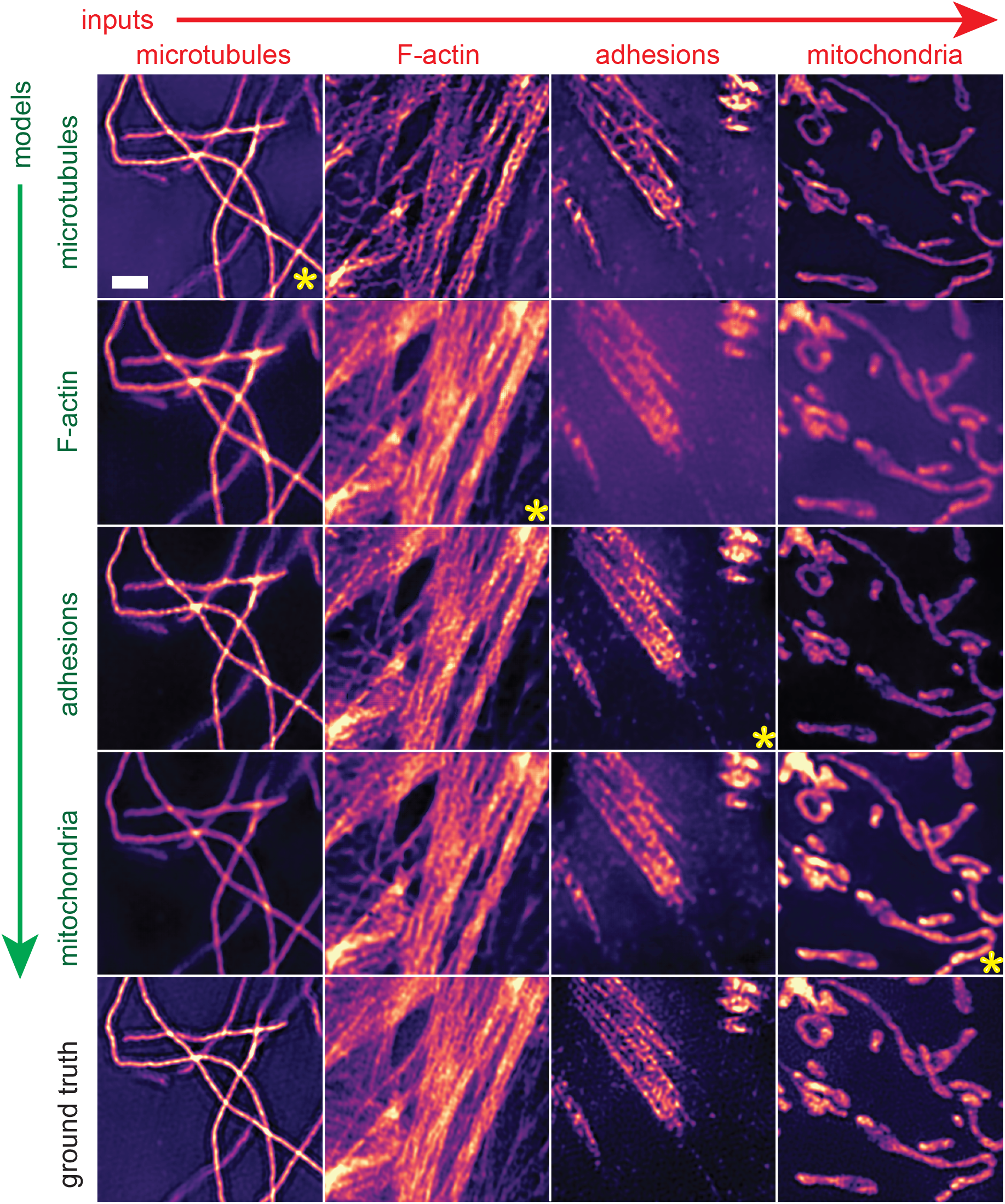
Apply a model trained on one structure to different structures. Each column shows input datasets of indicated structures. Each row shows the selected model pre-trained with the indicated structures and the last row shows the SIM reconstruction of the input dataset. The panels marked with a yellow asterisk were tested using the correct model. Scale bar: 1 μm.

**Supplementary Figure 9.**
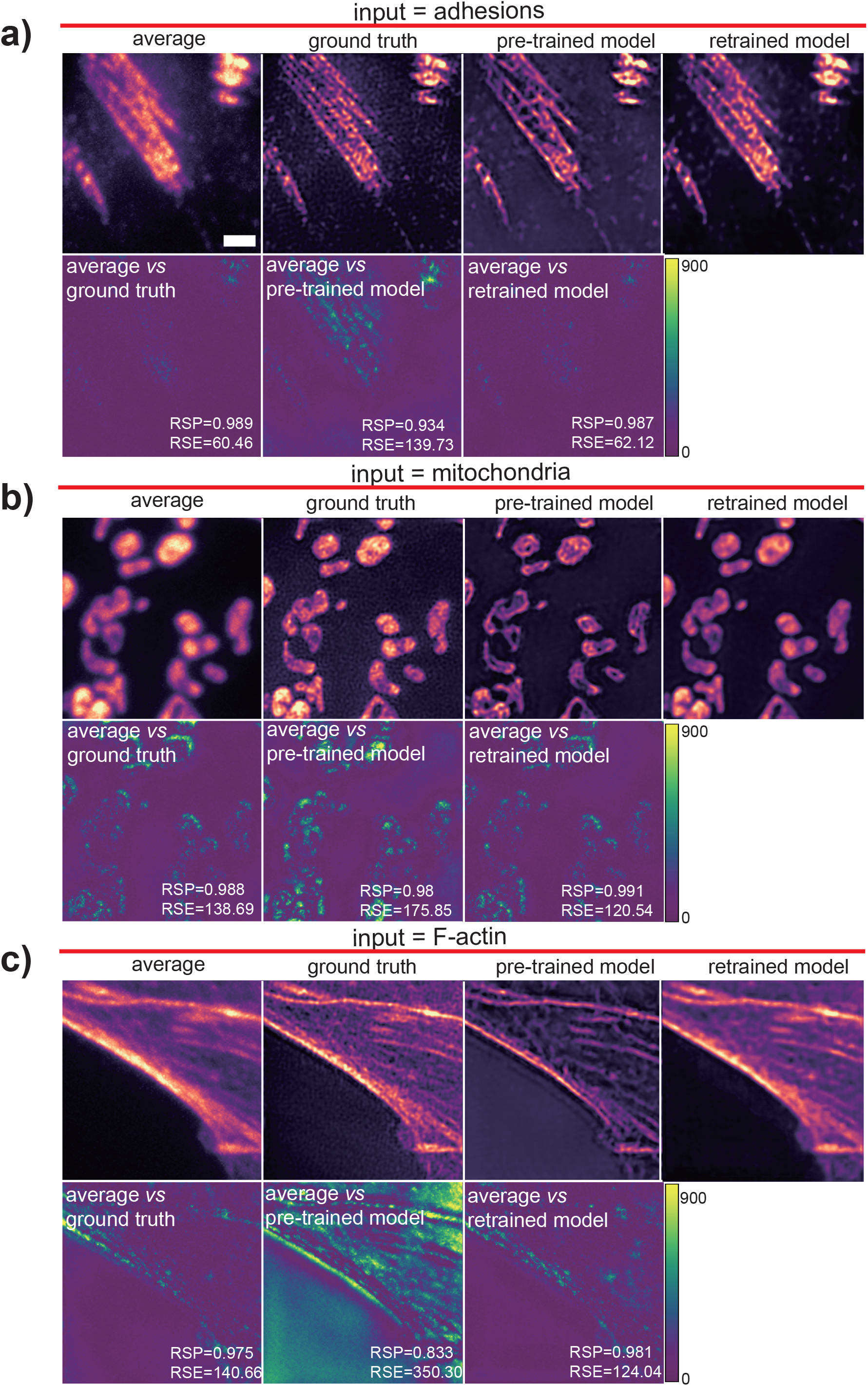
Transfer learning. To illustrate transfer learning, we made use of the network pre-trained on microtubules samples to initiate a new network and retrained it on other structures: adhesions (**a**), mitochondria (**b**) and F-actin (**c**). Fifteen SIM raw images were taken as the input. For each panel, top row: the average projection, the SIM reconstruction (ground truth), the output of the pre-trained model, and the output of re-trained model. The restoration error maps were estimated via SQUIRREL. The second row: SIM reconstruction, output of the pre-trained model and output of the retrained model. The value of RSE and RSP were calculated to quantify the restoration. Scale bar: 1 μm.

**Supplementary Videos 1. SR imaging of microtubules in living cells.** Shown are: the average projection of fifteen SIM raw images, SIM reconstruction, output of U-Net-SIM15 and output of U-Net-SIM3. Raw data was collected under normal light conditions (10% of 561 nm laser and 100 ms exposure time). Scale bar: 1 μm.

**Supplementary Videos 2. SR imaging of microtubules under low light conditions.** Shown are: the average projection of fifteen SIM raw images, SIM reconstruction, output of U-Net-SIM15 and output of scU-Net. Raw data was collected under low light conditions (1% of 561 nm laser and 5 ms exposure time). Scale bar: 1 μm.

**Supplementary Video 3. Dual-color imaging under low light conditions with scU-Net.** COS-7 cells were transfected with EMTB-3XmCherry overnight and stained with MitoTracker Green before imaging (**Methods**). Shown are: SIM reconstruction (left) and output of scU-Net (right). The data was collected using 2% of 488 nm laser and 1% of 561 nm laser with 50 ms exposure time. Scale bar: 1 μm.

**Supplementary Table 1.**
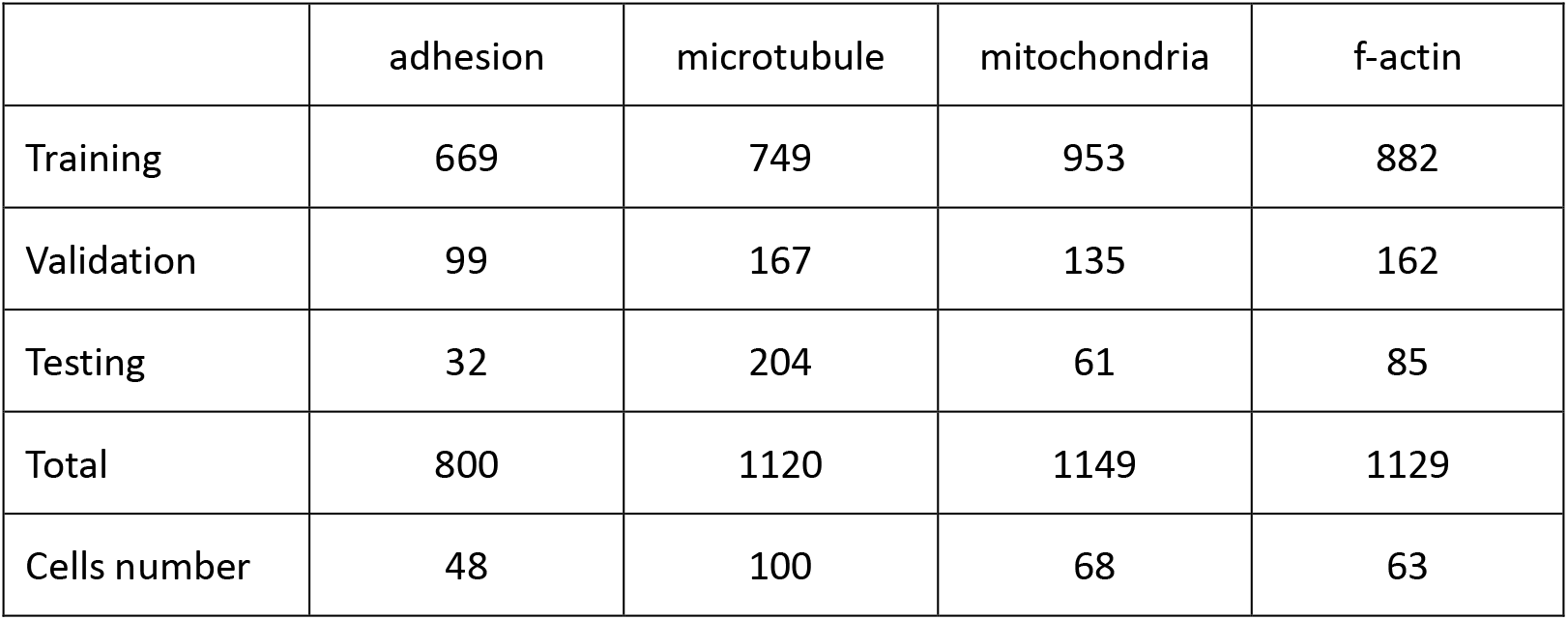
The size of the datasets for different structures.

|  | adhesion | microtubule | mitochondria | f-actin |
| --- | --- | --- | --- | --- |
| Training | 669 | 749 | 953 | 882 |
| Validation | 99 | 167 | 135 | 162 |
| Testing | 32 | 204 | 61 | 85 |
| Total | 800 | 1120 | 1149 | 1129 |
| Cells number | 48 | 100 | 68 | 63 |

**Supplementary Table 2.** Quantitative estimation of the performance of U-Net-SIM.

|  | Structures | Average | U-Net-SIM15 | U-Net-SIM3 |
| --- | --- | --- | --- | --- |
| PSNR | microtubules | $15.79 \pm 1.83$ | $35.75 \pm 5.44$ | $32.31 \pm 5.12$ |
| | adhesions | $17.92 \pm 1.92$ | $29.48 \pm 1.61$ | $24.84 \pm 1.89$ |
| | mitochondria | $20.32 \pm 1.95$ | $29.05 \pm 2.33$ | $27.29 \pm 2.39$ |
| | f-actin | $17.19 \pm 2.75$ | $21.59 \pm 4.82$ | $19.75 \pm 4.21$ |
| NRMSE | microtubules | $1.66 \pm 0.53$ | $0.19 \pm 0.10$ | $0.27 \pm 0.12$ |
| | adhesions | $1.06 \pm 0.30$ | $0.28 \pm 0.06$ | $0.47 \pm 0.10$ |
| | mitochondria | $0.63 \pm 0.14$ | $0.22 \pm 0.08$ | $0.28 \pm 0.09$ |
| | f-actin | $1.029 \pm 0.36$ | $0.70 \pm 0.46$ | $0.82 \pm 0.46$ |
| SSIM | microtubules | $0.19 \pm 0.08$ | $0.94 \pm 0.05$ | $0.91 \pm 0.07$ |
| | adhesions | $0.21 \pm 0.06$ | $0.77 \pm 0.06$ | $0.64 \pm 0.06$ |
| | mitochondria | $0.40 \pm 0.09$ | $0.83 \pm 0.05$ | $0.79 \pm 0.06$ |
| | f-actin | $0.21 \pm 0.10$ | $0.63 \pm 0.18$ | $0.53 \pm 0.15$ |

**Supplementary Table 3.** Quantitative estimation of the performance of U-Net-SIM15 and scU-Nets under low light condition.

|  | Structures | Average | Low light SIM | U-Net-SIM15 | scU-Nets |
| --- | --- | --- | --- | --- | --- |
| PSNR | microtubules | $15.93 \pm 1.82$ | $20.40 \pm 2.59$ | $26.93 \pm 4.30$ | $27.20 \pm 4.21$ |
| | adhesions | $18.15 \pm 1.98$ | $18.71 \pm 2.05$ | $23.09 \pm 2.18$ | $23.35 \pm 1.78$ |
| | mitochondria | $19.86 \pm 1.65$ | $20.32 \pm 2.54$ | $23.67 \pm 1.92$ | $23.75 \pm 1.94$ |
| | f-actin | $16.91 \pm 2.65$ | $11.09 \pm 7.64$ | $19.17 \pm 3.17$ | $18.84 \pm 3.82$ |
| NRMSE | microtubules | $1.66 \pm 0.51$ | $0.99 \pm 0.40$ | $0.49 \pm 0.17$ | $0.47 \pm 0.15$ |
| | adhesions | $1.07 \pm 0.30$ | $0.83 \pm 0.26$ | $0.57 \pm 0.13$ | $0.53 \pm 0.12$ |
| | mitochondria | $0.68 \pm 0.15$ | $0.60 \pm 0.16$ | $0.40 \pm 0.09$ | $0.40 \pm 0.09$ |
| | f-actin | $1.07 \pm 0.35$ | $2.21 \pm 1.38$ | $0.84 \pm 0.36$ | $0.88 \pm 0.47$ |
| SSIM | microtubules | $0.17 \pm 0.07$ | $0.43 \pm 0.14$ | $0.83 \pm 0.12$ | $0.83 \pm 0.12$ |
| | adhesions | $0.20 \pm 0.05$ | $0.33 \pm 0.08$ | $0.55 \pm 0.09$ | $0.55 \pm 0.08$ |
| | mitochondria | $0.33 \pm 0.09$ | $0.50 \pm 0.11$ | $0.71 \pm 0.07$ | $0.69 \pm 0.08$ |
| | f-actin | $0.20 \pm 0.09$ | $0.38 \pm 0.14$ | $0.42 \pm 0.16$ | $0.44 \pm 0.14$ |

